# Prognostic value, signal interaction network, and immune infiltration characteristics of MET gene expression in gastric cancer analyzed by multi-database bioinformatics

**DOI:** 10.64898/2026.09.23.753715

**Authors:** Zhengjie Zhang, Xiuzhen Wei

**Affiliations:** Department of Gastroenterology, Wuwei Liangzhou Hospital, China,733000; Department of Oncology, Wuwei Liangwhou Hospital, China,,733000

**Author notes:** Corresponding author: Xiuzhen Wei.

**Keywords:** Gastric Carcinoma, Hepatocyte Growth Factor Receptor (MET), Bioinformatics, Prognosis, Immune Infiltration

## Abstract

**Objective:** Based on the bioinformatics method of multi-database integration, this study systematically analyzes the expression characteristics, clinical pathological correlation, prognostic value, potential molecular mechanisms, and immune infiltration patterns of hepatocyte growth factor receptor (MET) in gastric cancer.

**Methods:** The UALCAN and GEPIA databases were employed to examine the differential expression of MET between gastric cancer and normal gastric mucosal tissues, as well as its associations with clinicopathological features. Kaplan-Meier Plotter was utilized to evaluate the impact of MET expression on overall survival (OS) and progression-free survival (PFS). Protein-protein interaction (PPI) network was constructed via LinkedOmics, followed by Gene Ontology (GO) functional annotation and Kyoto Encyclopedia of Genes and Genomes (KEGG) pathway enrichment analysis of co-expressed genes. Four algorithms, TIMER, CIBERSORT, EPIC, and MCPcounter, were used to cross evaluate the correlation between MET expression and immune cell infiltration; Further validate the cell type specific expression of MET using the gastric cancer single-cell sequencing queues (GSE134520, GSE167297) built into the TISCH database.

**Results:** MET expression was significantly elevated in gastric cancer tissues compared with normal gastric mucosa (*P* < .05), and its expression level was significantly correlated with tumor grade and TNM stage. Patients with high MET expression exhibited significantly poorer OS and PFS than those with low MET expression (*P* < .05). The PPI network revealed that MET could interact with 20 key proteins, including EGFR, ERBB2, HGF, STAT3, and GRB2 etc. GO enrichment analysis suggests that differentially expressed genes are significantly enriched in functions such as the ERBB signaling pathway, cadherin binding, and DNA repair complexes; KEGG enrichment analysis showed that MET related genes were significantly enriched in pathways such as homologous recombination, nuclear cytoplasmic transport, mismatch repair, and oxidative phosphorylation. Immune infiltration analysis showed that the negative association between MET and B cells infiltration has cross algorithm robustness, while the association with neutrophils, CD8^+^ T cells, CD4^+^ T cells, and macrophages exhibits algorithmic heterogeneity or insignificance; There is no significant correlation between MET and common immune checkpoint molecules such as PD-1, PD-L1, CTLA4, etc. Single cell validation further confirmed that MET is mainly enriched in malignant epithelial cells and endothelial cells, and is almost not expressed in immune cells.

**Conclusions:** Multidimensional bioinformatic analyses demonstrate that elevated MET expression serves as an independent risk factor for unfavorable prognosis in gastric cancer. MET may mediate dual drug resistance in gastric cancer via crosstalk with multiple signaling molecules (including EGFR, ERBB2, HGF, STAT3 and GRB2) and dysregulation of the homologous recombination repair pathway. Results from multiple-algorithm immune infiltration analysis, single-cell dataset analysis and immune checkpoint correlation analysis indicate that MET exerts only modest direct regulatory effects on the gastric cancer immune microenvironment. This exploratory study offers systematic bioinformatic evidence supporting MET as a candidate prognostic biomarker and potential therapeutic target for gastric cancer. Further functional experiments and prospective cohort studies are required to validate its molecular mechanisms and clinical utility.

---

According to cancer statistics released in the United States, an estimated 31,510 new cases of gastric cancer and 10,740 deaths are projected for 2026[1]. Despite a decline in incidence, gastric cancer remains a major threat to human health[2]. Current, the treatment strategy for gastric cancer is gradually expanding from traditional chemotherapy to molecular targeted therapy and immunotherapy, treatment modalities for advanced gastric cancer primarily include chemotherapy, targeted therapy, and immunotherapy[3-5]. Approved targeted agents include trastuzumab targeting human epidermal growth factor receptor 2 (HER-2)[6] and zolbetuximab targeting Claudin18.2[7]. However, only a limited subset of patients benefit from these agents, and most eventually develop drug resistance[8]. Therefore, identifying more reliable prognostic markers and effective therapeutic targets is of substantial clinical significance for improving the diagnosis, treatment, and survival outcomes of gastric cancer patients.

Hepatocyte growth factor receptor (MET) is a receptor tyrosine kinase encoded by the oncogene MET. Abnormal activation of MET can promote tumor cell proliferation, migration, invasion, and angiogenesis through multiple signaling pathways such as PI3K-AKT, RAS-MAPK, and STAT3[9].In gastric cancer, the incidence of MET overexpression is as high as 50% - 65%[10], suggesting that it may play a key role in the occurrence and development of gastric cancer. As early as 1992, relevant literature reported the association between gastric cancer and MET inhibitors[11]. Until recently, research on the mechanism of action and targeted therapeutic value of MET in gastric cancer has continued to deepen[12]. However, previous studies have mostly focused on the pro proliferative and pro metastatic functions of MET, and there is still a lack of systematic interpretation of its role in treating drug resistance and its interaction with the tumor immune microenvironment. Moreover, most bioinformatics studies rely on a single database or analysis method, and the robustness of the conclusions needs further verification.

This study systematically describes the expression characteristics, clinical pathological correlation, and prognostic value of MET in gastric cancer through bioinformatics analysis integrating multiple databases; Utilizing protein interaction networks and functional enrichment analysis to explore potential molecular regulatory mechanisms, with a focus on exploring the role of MET in targeted drug resistance and chemotherapy resistance; And using four immune deconvolution algorithms combined with single-cell external queue data, the relationship between MET and tumor immune microenvironment is analyzed from multiple dimensions. The aim of this study is to provide more systematic bioinformatics evidence for MET as a prognostic biomarker and potential therapeutic target for gastric cancer, and to provide new theoretical references for optimizing the combination therapy strategy for gastric cancer with high MET expression.

## 1. Data and Methods

### 1.1 Data Sources and Differential Expression Analysis

MET expression data across cancers were obtained from the Gene Expression Profiling Interactive Analysis (GEPIA) database (version 2, http://gepia.cancer-pku.cn/), which integrates data from The Cancer Genome Atlas (TCGA) and the Genotype-Tissue Expression (GTEx) project. A total of 408 gastric cancer tissue samples, 211 normal gastric tissue samples, and clinical data from 629 patients were extracted.

For validation, MET expression data were also retrieved from the UALCAN database (http://ualcan.path.uab.edu/), including 415 gastric cancer tissue samples, 34 normal gastric tissue samples, and clinical data from 449 patients.

Differential expression of MET between gastric cancer and normal tissues was analyzed using the Wilcoxon rank-sum test. Patients were stratified according to clinical characteristics, including sex, age, tumor grade, TNM stage, H. pylori (HP) infection and TP53 mutation, to assess the association between MET expression levels and clinicopathological features.

### 1.2 Survival Analysis

Survival analysis was performed using the Kaplan-Meier Plotter database (http://kmplot.com/analysis/). This database integrates transcriptome data and clinical follow-up information from multiple public databases, including Gene Expression Omnibus (GEO), European Genome phenome Archive (EGA), and The Cancer Genome Atlas (TCGA), for gastric cancer. This database contains overall survival (OS) data for 881 patients and progression free survival (PFS) data for 645 patients. Patients were divided into high--MET and low-MET expression groups based on the optimal cutoff value determined by the Kaplan-Meier Plotter algorithm, which maximizes the log-rank test statistic. The correlation between MET expression levels and patient OS and PFS was evaluated using Kaplan-Meier survival curves and the log-rank test. Hazard ratios (HRs) with 95% confidence intervals (CIs) were calculated.

### 1.3 Co-expression and Functional Enrichment Analysis

Genes co-expressed with MET in gastric cancer were identified using the LinkedOmics database (http://www.linkedomics.org/). Pearson correlation coefficients were calculated, and genes with a false discovery rate (FDR) < 0.05 were considered significantly correlated with MET.

Gene Ontology (GO) functional annotation and Kyoto Encyclopedia of Genes and Genomes (KEGG) pathway enrichment analysis were performed on the co-expressed gene set using the LinkedOmics analysis module. An FDR-adjusted P-value < .05 was set as the threshold for statistical significance.

### 1.4 Protein-Protein Interaction Network Analysis

Protein-protein interaction (PPI) prediction for MET in gastric cancer was performed using the STRING database (https://string-db.org/), restricted to Homo sapiens. Interacting proteins with a combined confidence score > 0.7 (high confidence) were retrieved, and the top 20 interactors ranked by score were selected for subsequent network analysis. Visualization of the PPI network was conducted using the bioinformatics platform (https://www.bioinformatics.com.cn).

### 1.5 Immune Infiltration Analysis

#### 1.5.1 TIMER Database Analysis

The correlation between MET expression and immune cell infiltration in gastric cancer was analyzed using the Tumor IMmune Estimation Resource (TIMER) database (https://cistrome.shinyapps.io/timer/). Spearman correlation analysis was performed to assess the relationship between MET expression and the infiltration levels of immune cells, including B cells, CD4^+^ T cells, CD8^+^ T cells, macrophages, and neutrophils.

#### 1.5.2 Multi algorithm cross validation

To verify the robustness of TIMER results, three independent deconvolution algorithms, CIBERSORT, EPIC, and MCPcounter, were further used to re estimate the infiltration abundance of the immune cell subpopulations based on gene expression profile data from the TCGA-STAD cohort. Pearson correlation analysis was used to evaluate the association between MET expression and immune scores of various algorithms.

#### 1.5.3 Single cell transcriptome external queue validation

Clarify the cell type specific expression of MET at single-cell resolution using the TISCH database (http://tisch.comp-genomics.org/) built in two independent single-cell RNA sequencing datasets for gastric cancer (GSE134520 and GSE167297). Extract standardized expression values (log (TPM/10+1)) of MET in various cell types and generate cell type specific expression heatmaps.

#### 1.5.4 Correlation analysis of immune checkpoint genes

Using the GEPIA2 database based on the TCGA-STAD gastric cancer cohort, analyze the expression correlation between MET and six classic immune checkpoint genes (PDCD1 (PD-1), PD-L1, CTLA4, TIGIT, LAG3, HAVCR2). All gene expression values were standardized using log₂ (TPM), and Pearson correlation analysis was used to calculate the correlation coefficient R and corresponding *P* value. A scatter plot was plotted to visualize the correlation between the two.

### 1.6 Statistical Analysis

All statistical analyses were performed using SPSS software (version 25.0). Between-group comparisons were conducted using the independent t-test for normally distributed data or the Mann-Whitney U test for non-normally distributed data. Associations between MET expression and categorical clinical variables were assessed using the Pearson χ² test (chi-square test). Correlation analyses were performed using Pearson or Spearman rank correlation, as appropriate. A two-sided P-value < .05 was considered statistically significant, except for enrichment analyses where FDR adjustment was applied.

### 1.7 Ethical Approval

This study utilized publicly available databases (GEPIA, UALCAN). No human participants or animal subjects were involved. Therefore, ethical approval was not required.

## 2 Results

### 2.1 Expression of MET in Gastric Cancer

Pan-cancer analysis based on the GEPIA database revealed that MET expression levels in gastric cancer (STAD), cervical squamous cell carcinoma (CESC), colon adenocarcinoma (COAD), esophageal cancer (ESCA), and renal cancer (KICH) were significantly higher than those in corresponding normal tissues ( *P* < .05). In contrast, MET expression was low in breast cancer (BRCA), acute myeloid leukemia (LAML), and low-grade glioma (LGG), suggesting that MET expression exhibits cancer-type specificity (Figure 1).We further analyzed MET expression differences using data from the UALCAN database (415 gastric cancer tissues *vs* 34 normal gastric mucosal tissues) and the GEPIA database (408 gastric cancer tissues *vs* 211 normal gastric mucosal tissues). Both databases consistently demonstrated that MET expression levels were significantly elevated in gastric cancer tissues compared with normal gastric mucosal tissues ( *P* < .05) (Table 1).

**Fig. 1.**
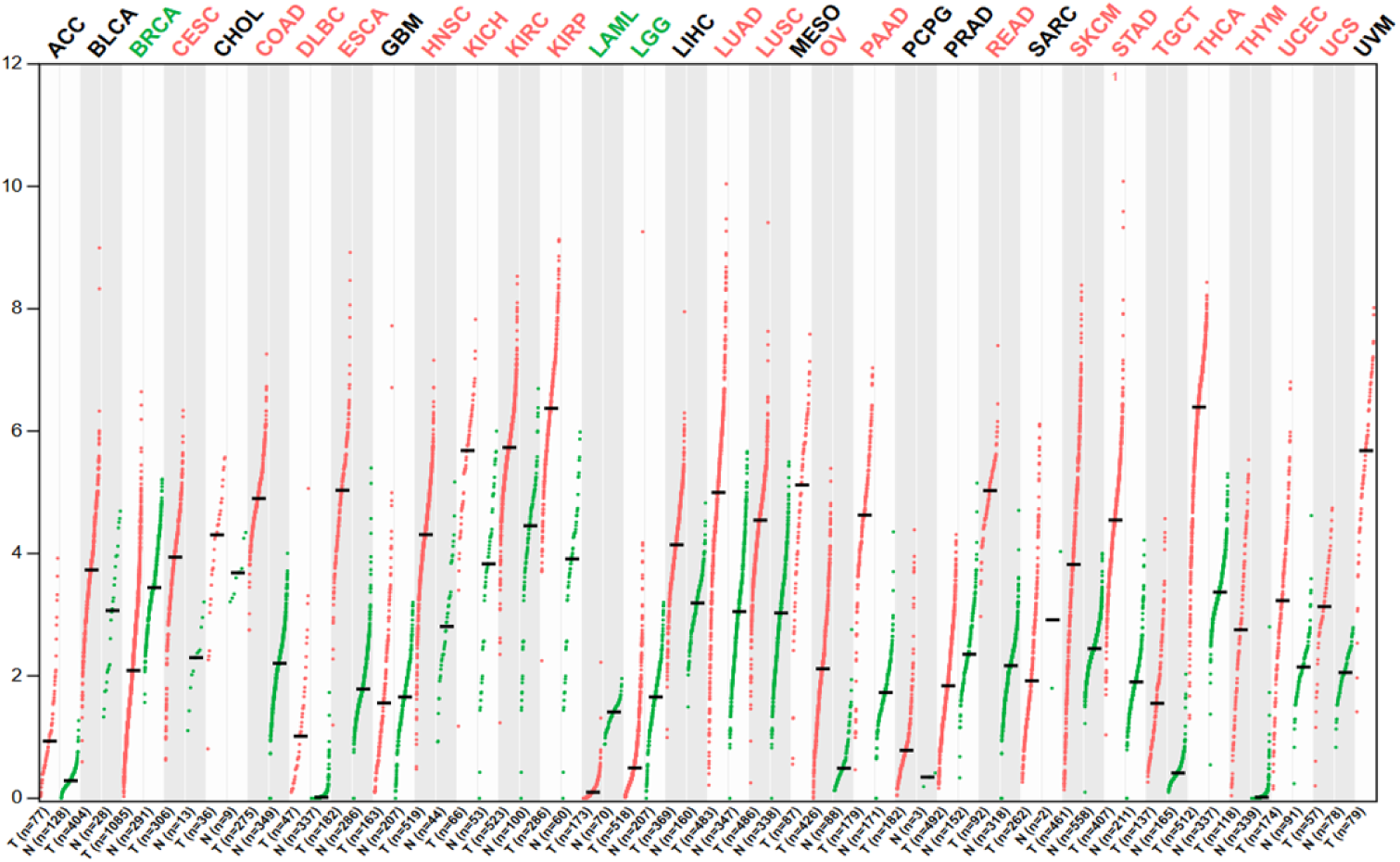
Expression of MET in Cancers Based on GEPIA Database (red represents high expression, green represents low expression)

**Tab. 1:**
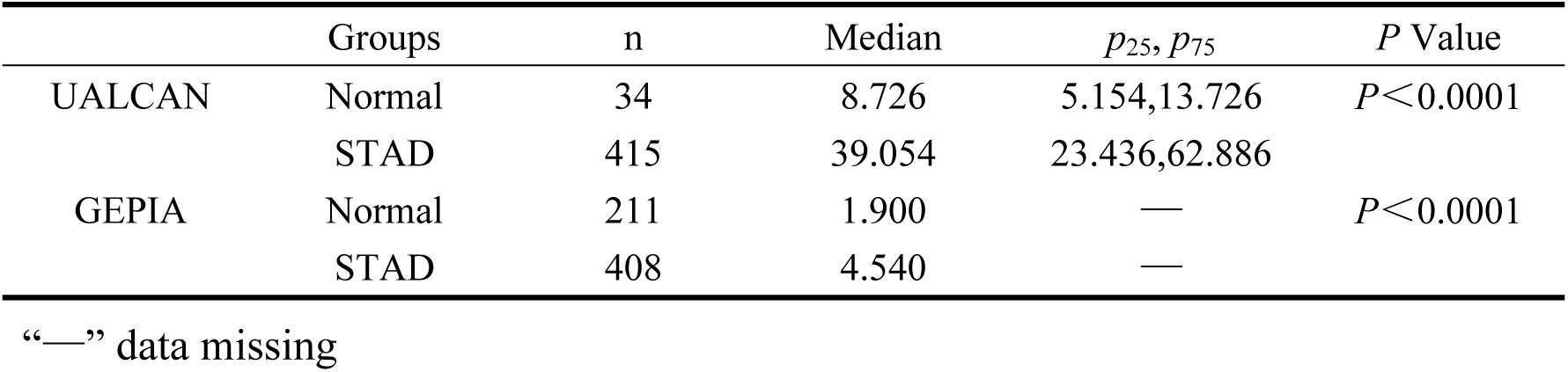
Expression of MET in Gastric Cancer Tissue (STAD) and Normal Tissue [M (*p*25, *p*75)]

| | Groups | n | Median | $p_{25}, p_{75}$ | $P$ Value |
| --- | --- | --- | --- | --- | --- |
| UALCAN | Normal | 34 | 8.726 | 5.154,13.726 | $P < 0.0001$ |
|  | STAD | 415 | 39.054 | 23.436,62.886 |  |
| GEPIA | Normal | 211 | 1.900 | — | $P < 0.0001$ |
|  | STAD | 408 | 4.540 | — |  |

“—” data missing

### 2.2 Correlation Between MET Expression and Clinicopathological Characteristics

Clinical data of 415 gastric cancer patients were retrieved from the UALCAN database to analyze the correlation between MET expression and clinicopathological characteristics.

MET expression levels were significantly higher in gastric cancer patients than in the normal population across all analyzed subgroups, including stratification by sex, age, tumor grade, lymph node metastasis, tumor stage, HP infection, and TP53 mutation. With the exception of the subgroups aged 21–40 years (*P* = .165) and clinical stage 4 (*P* = .060), the differences in MET expression between gastric cancer patients and normal controls were statistically significant in all other subgroups (*P* < .050) (Table 1).

In gastric cancer patients, MET expression levels in patients with tumor grade 1 were lower than those in grades 2 and 3, with a statistically significant difference observed between grade 1 and grade 3 (*P* < .001); However, no significant differences were detected between grades 1 and 2, or between grades 2 and 3. Regarding TNM stage, MET expression levels in TNM stage 3 patients were significantly higher than those in stage 1 (*P* = .010) and stage 2 (*P* = .043), whereas no significant differences were observed between stages 1 and 2, stages 1 and 4, or stages 2 and 4. No significant differences in MET expression levels were detected across subgroups stratified by sex, age, HP, or TP53 mutation ( all *P* > .050). The detailed results are presented in Table 2 and Figure 2.

**Fig. 2.**
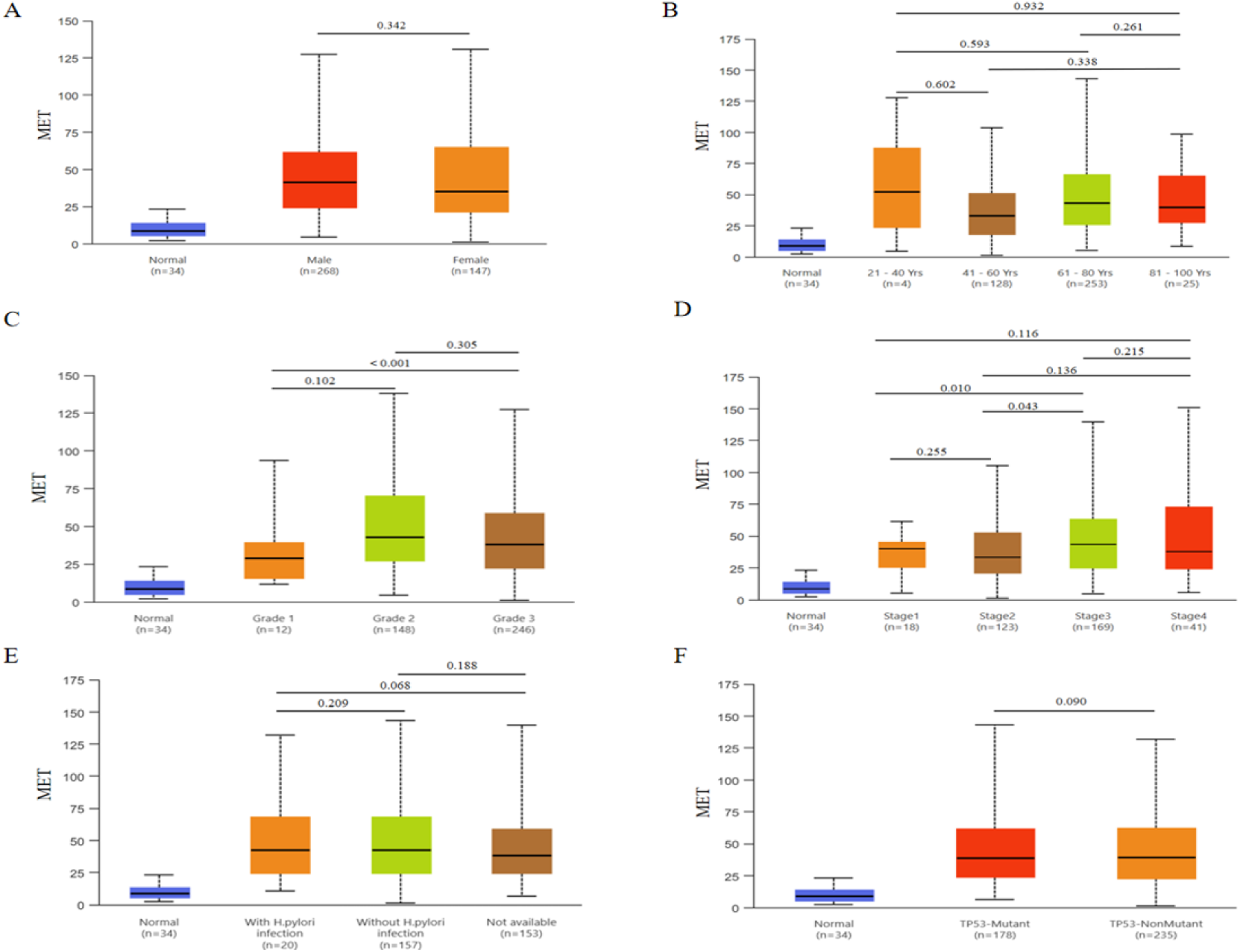
MET expression in gastric cancer patients A: Correlation between MET and gender; B: Correlation between MET and age; C: Correlation between MET and tumor grade; D: Correlation between MET and TNM stage; E: Correlation between MET and HP; F: Correlation between MET and TP53.

**Tab. 2.** Correlation between MET expression and clinical pathological characteristics of gastric cancer patients.

| Characteristics | n | M ($p_{25}, p_{75}$) | P Value |
| --- | --- | --- | --- |
| Normal | 34 | 8.785 (5.305,13.436) |  |
| Gender |  |  |  |
| Male | 268 | 41.179 (24.120, 61.186) | <0.001 |
| Female | 147 | 34.950 (21.408, 64.671) | 0.002 |
| Age |  |  |  |
| 21-40 Yrs | 4 | 52.255 (23.878, 87.570) | 0.165 |
| 41-60 Yrs | 128 | 32.938 (18.041, 50.862) | <0.001 |
| 61-80 Yrs | 253 | 42.946 (25.951, 65.785) | <0.001 |
| 81-100 Yrs | 25 | 39.665 (27.894, 65.112) | <0.001 |
| Grade |  |  |  |
| 1 | 12 | 28.908 (15.749, 39.321) | 0.009 |
| 2 | 148 | 42.982 (27.145, 69.840) | <0.001 |
| 3 | 246 | 37.892 (22.528, 58.312) | <0.001 |
| N |  |  |  |
| N0 | 123 | 38.740 (23.206, 62.589) | <0.001 |
| N1 | 112 | 36.155 (21.735, 59.187) | <0.001 |
| N2 | 79 | 41.337 (25.257, 57.811) | 0.002 |
| N3 | 82 | 39.260 (24.095, 66.052) | 0.015 |
| Stages |  |  |  |
| 1 | 18 | 40.288 (25.364, 45.367) | <0.001 |
| 2 | 123 | 33.086 (20.694, 52.591) | <0.001 |
| 3 | 169 | 43.371 (24.883, 62.911) | <0.001 |
| 4 | 41 | 37.892 (24.379, 72.637) | 0.06 |
| HP |  |  |  |
| With HP | 20 | 42.453 (24.346, 68.174) | <0.001 |
| Without HP | 157 | 42.604 (24.243, 67.870) | <0.001 |
| Not available | 153 | 38.407 (24.428, 58.842) | <0.001 |
| TP53 |  |  |  |
| TP53-Mutant | 178 | 38.815 (23.722, 61.370) | <0.001 |
| TP53-NonMutant | 235 | 39.448 (22.807, 62.217) | <0.001 |

### 2.3 Correlation Between MET Expression and Prognosis in Gastric Cancer

Kaplan-Meier survival analysis using the Kaplan-Meier Plotter database showed that gastric cancer patients with high MET expression had significantly worse OS and PFS compared with those with low MET expression. The HR for OS was 1.58 (95% CI: 1.33–1.89), and the HR for PFS was 1.59 (95% CI: 1.30–1.95). These results indicate that MET overexpression is associated with poor prognosis in gastric cancer patients (Figure 3).

**Fig. 3:**
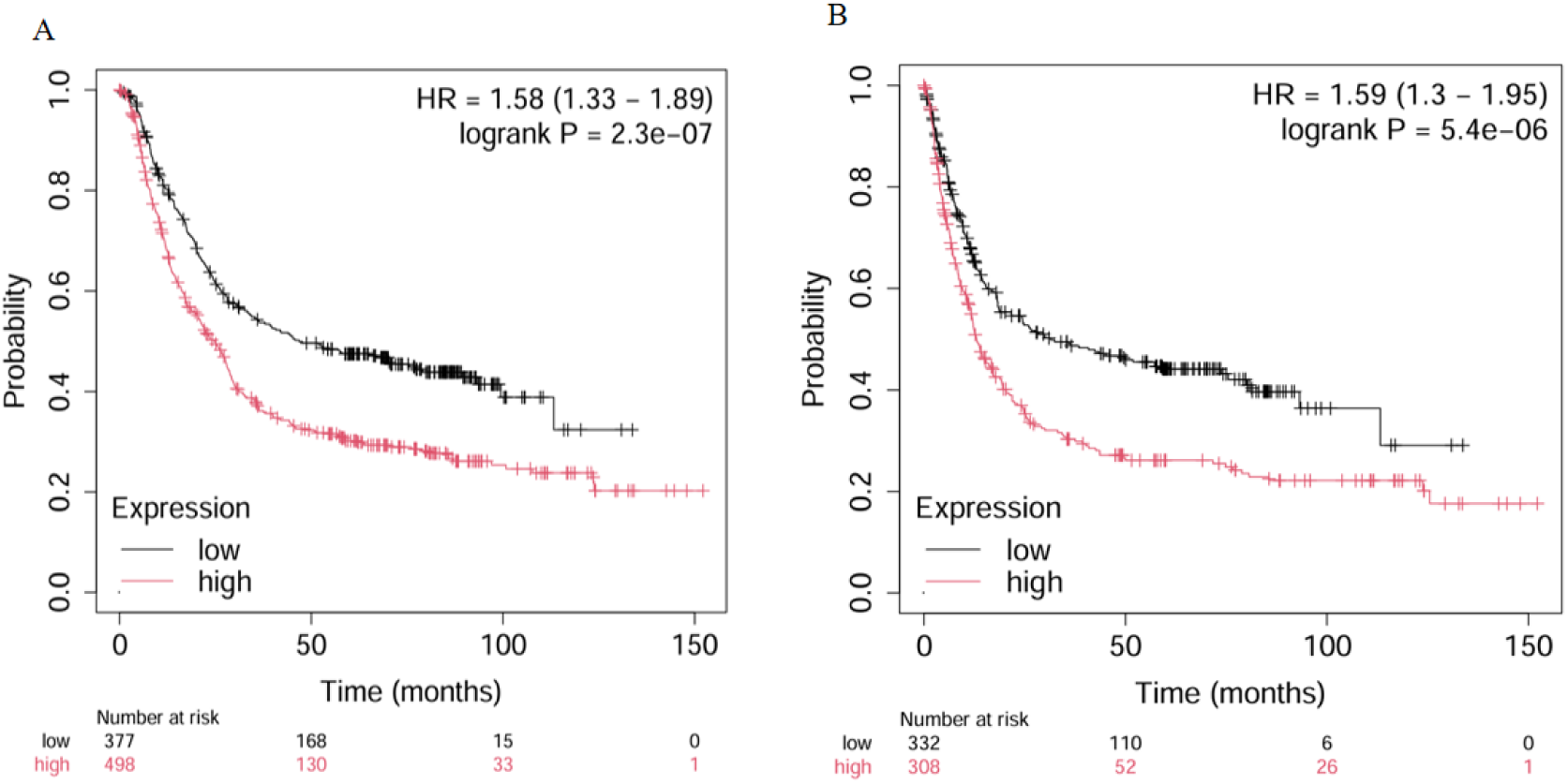
Prognostic value of MET in gastric cancer. A: Relationship between MET and gastric cancer OS; B: The relationship between MET and gastric cancer PFS

### 2.4 MET Related Genes in Gastric Cancer

Using the LinkedOmics database to analyze 415 gastric cancer tissue samples, we identified 50 genes significantly correlated with MET expression (absolute Pearson correlation > 0.3, FDR < 0.05). Among these, genes positively correlated with MET expression included CAPZA2, ST7, TES, ITGA2, and LMO7, while genes negatively correlated with MET expression included GKAP1, OAZ2, C3orf18, RNF122, and IL11RA (Figures 4 A and 4B).

**Fig. 4.**
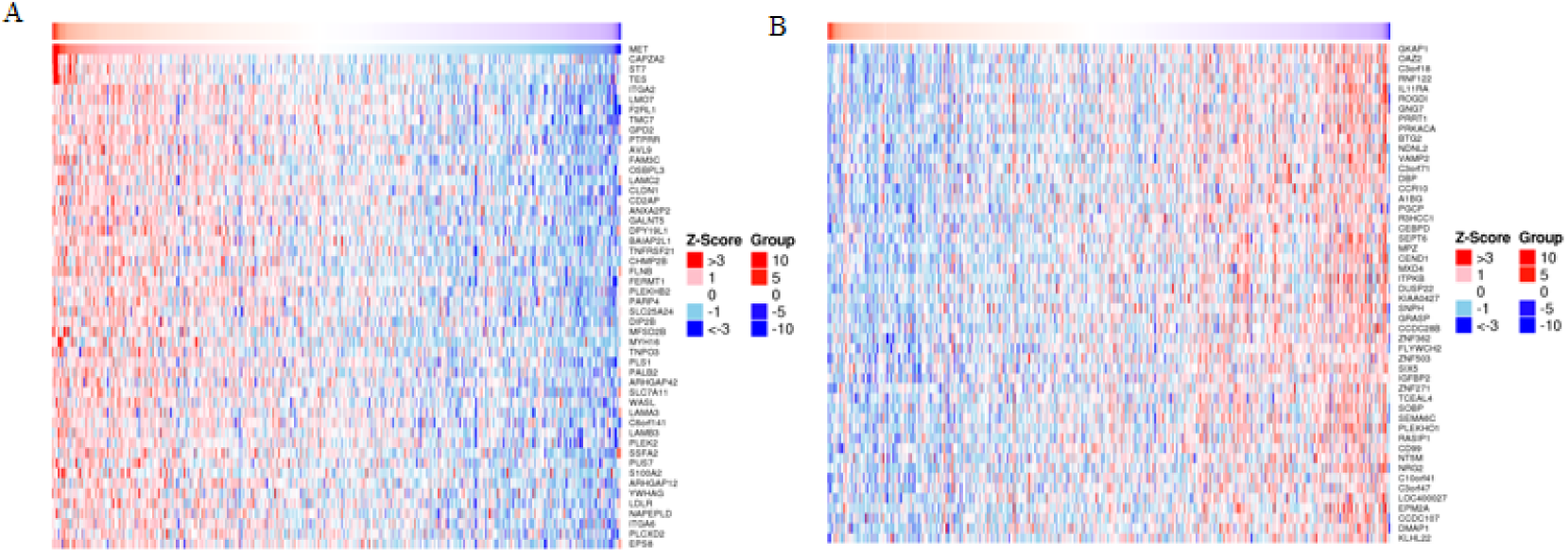
MET related genes in gastric cancer samples. A: MET positively correlated genes; B: MET negatively correlated genes

### 2.5 Functional Enrichment Analysis of MET

To explore the potential molecular mechanism of MET affecting the progression of gastric cancer, we conducted GO and KEGG enrichment analysis on differentially expressed genes between MET high and low expression groups. GO biological process (BP) enrichment showed that differentially expressed genes were significantly enriched in biological processes such as the ERBB signaling pathway and integrin mediated cell adhesion (Figure 5A); The enrichment of GO cell components (CC) suggests significant enrichment of DNA repair complexes and chromatin related structures (Figure 5B); The GO molecular function (MF) enrichment results showed a significant enrichment of cadherin binding function, while oxidoreductase related functions only showed an enrichment trend, with no statistical significance (FDR > 0.05, Figure 5C).

**Fig. 5.**
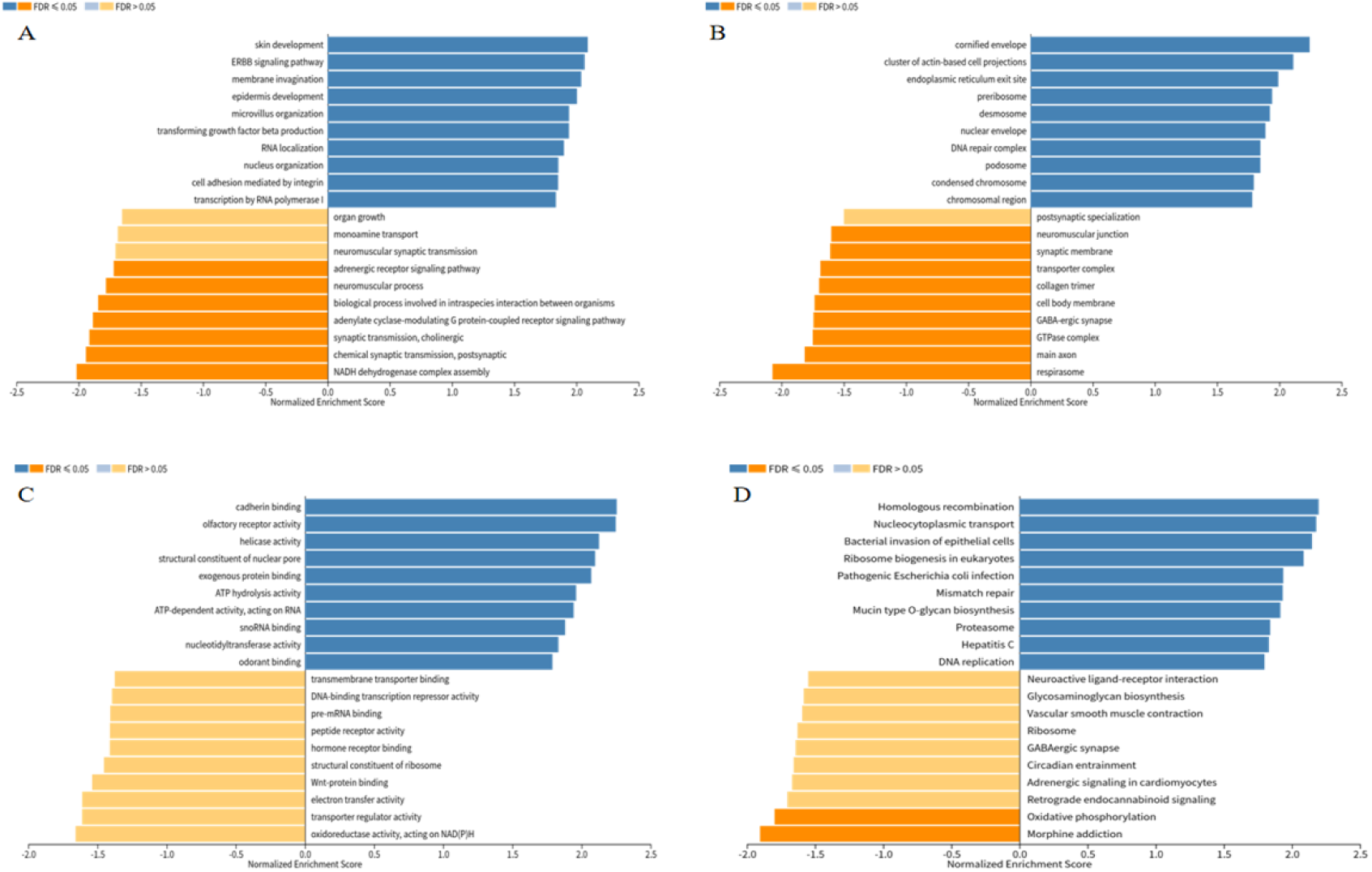
Functional enrichment analysis of MET. A: Biological progress (BP); B: Cellular component (CC); C: Molecular function (MF); D: KEGG pathway

KEGG enrichment analysis further revealed that MET related differentially expressed genes were significantly enriched in pathways such as homologous recombination, nuclear cytoplasmic transport, mismatch repair, and oxidative phosphorylation (Figure 5D). The significant enrichment of homologous recombination and mismatch repair pathways provides key clues for MET mediated chemotherapy resistance, while the enrichment of ERBB signaling pathway suggests that RTK cross activation is the molecular basis of targeted resistance. These results collectively indicate that the oncogenic role of MET in gastric cancer involves a multidimensional and multi-level molecular regulatory network.

### 2.6 PPI Network of MET

To systematically depict the signaling network of MET in gastric cancer, we constructed a MET protein interaction network using the STRING database and identified 20 protein molecules that significantly interact with MET, including EGFR, ERBB2, HGF, STAT3, and GRB2 (Figure 6). At the membrane level, MET directly interacts with RTK family members EGFR and ERBB2, suggesting that MET may form a cross activated membrane receptor signaling network with other RTKs. This may be an important molecular mechanism for the limited clinical efficacy of single MET inhibitors or EGFR inhibitors; At the cytoplasmic level, MET is directly associated with HGF and GRB2, forming the classical downstream signaling core axis of MET; At the nuclear level, MET interacts with the transcription factor STAT3, indicating that MET signaling can penetrate deep into the nucleus and exert biological functions by regulating target gene transcription.

**Fig. 6.**
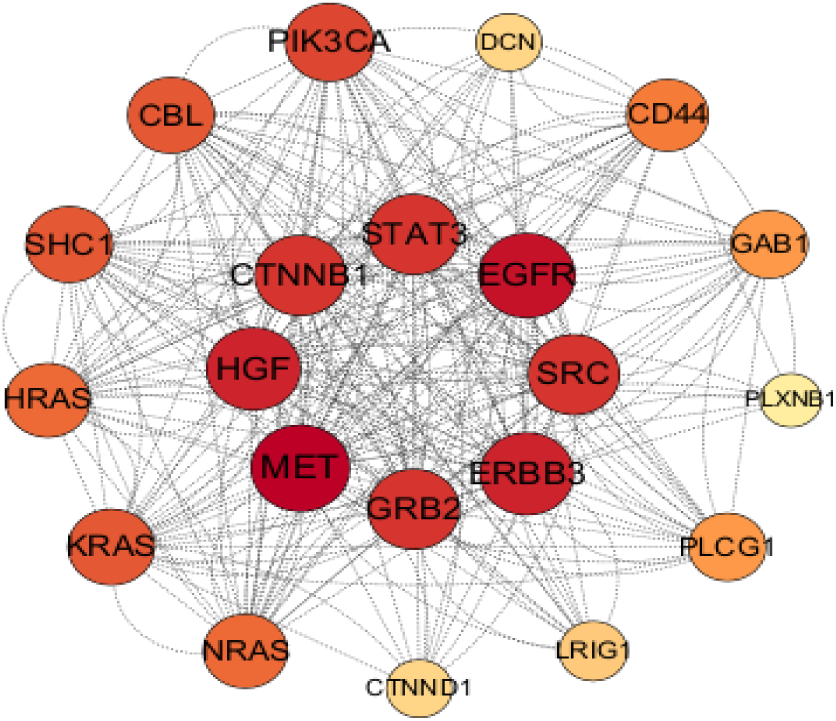
PPI network diagram of MET gene

### 2.7 Multi algorithm cross validation of MET expression and immune cell infiltration

We first explore the relationship between MET expression and the immune microenvironment of gastric cancer using the TIMER database. The results showed no significant correlation between MET expression and tumor purity (Rho=0.006, *P*=.246), ruling out the confounding effect of tumor cell content on immune infiltration assessment. In the immune cell subpopulation, MET expression was negatively correlated with B cells (Rho=-0.127, *P*=.014) and CD4⁺ T cells (Rho=-0.118, *P*= .022), weakly positively correlated with CD8⁺ T cells (Rho=0.103, *P*= .047), and not significantly correlated with macrophages and neutrophils (Figure 7).

**Fig. 7.**
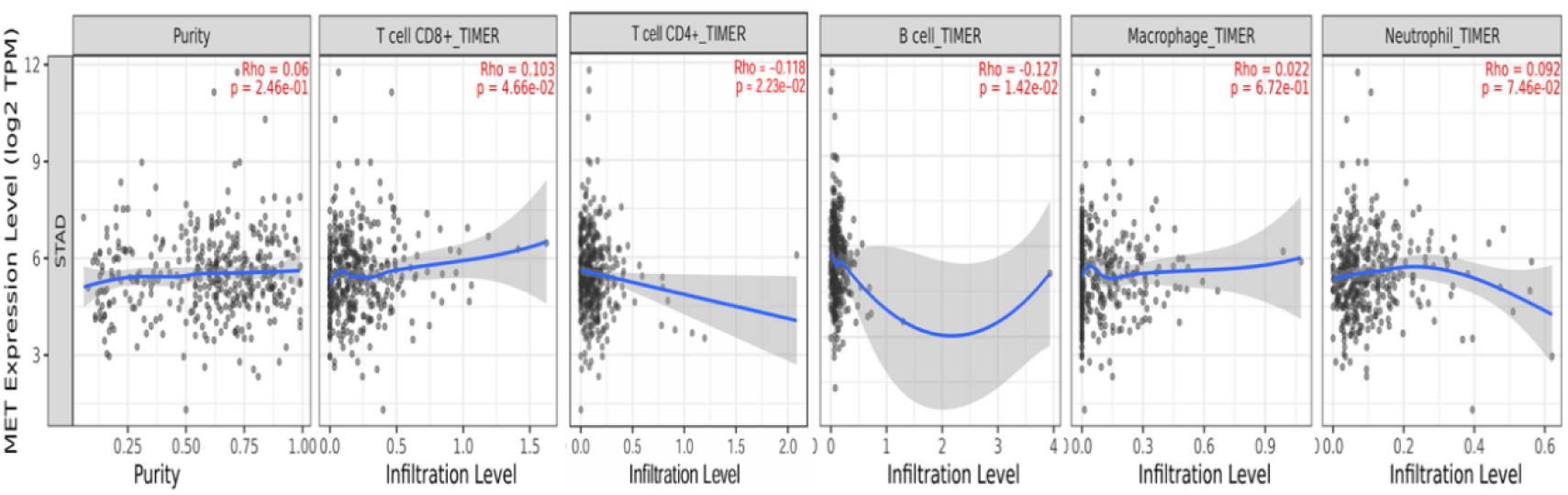
Exploratory screening of MET expression and immune cell infiltration

To verify the robustness of the above results, we further conducted cross validation using three independent algorithms: CIBERSORT, EPIC, and MCPcounter. The results showed that the negative correlation of B cells had the highest cross algorithm consistency: all three algorithms confirmed a significant negative correlation between MET expression and B cells infiltration (all *P* < .01). Neutrophils showed a consistent weak positive correlation in both CIBERSORT and MCPcounter (both *P* < .05). There is significant heterogeneity in the results of CD4⁺ T cells among different algorithms (EPIC shows positive correlation, CIBERSORT shows no significant correlation). The weak positive correlation between MET and CD8⁺ T cells suggested in TIMER analysis was not detected by CIBERSORT, EPIC, and MCPcounter algorithms, and the absolute values of the correlation coefficients were all less than 0.10 (all *P*> .05), indicating that this correlation does not have cross platform robustness. For macrophages, there was no significant correlation between CIBERSORT and EPIC, only weak positive correlation was detected by MCPcounter (r=0.153, *P*=.003), and there was significant heterogeneity in the results of different algorithms, with no stable and reliable evidence of association (Table 3). The negative correlation between MET and B cells remained consistent and significant in all four analysis methods.

**Tab. 3.** Correlation analysis between MET gene expression levels and multiple algorithms for estimating tumor immune cell infiltration abundance.

| Immune cell types | CIBERSORT($r/P$) | EPIC($r/P$) | MCPcounter($r/P$) |
| --- | --- | --- | --- |
| CD8<sup>+</sup> T cell | -0.075 / 0.149 | -0.056 / 0.283 | -0.041 / 0.432 |
| CD4<sup>+</sup> T cell | 0.045 / 0.388 | 0.143 / 0.006 | - |
| B cells | -0.219 / <0.001 | -0.228 / <0.001 | -0.156 / 0.002 |
| Macrophage | 0.075 / 0.146 | 0.025 / 0.634 | 0.153 / 0.003 |
| Neutrophil | 0.133 / 0.010 | - | 0.157 / 0.002 |

“-” missing data

### 2.8 Cell type specific expression of MET gene in single-cell transcriptome of gastric cancer

To address the limitations of relying solely on a large number of transcriptome deconvolution algorithms for immune infiltration analysis, we further supplemented single-cell RNA seq data from two independent external cancer cohorts (GSE134520 and GSE167297) to characterize the cell type specific expression patterns of MET. As shown in the heatmap results of Figure 8, MET is almost not expressed in CD8⁺ T cells, B cells, dendritic cells, and other immune groups. Its expression is mainly enriched in malignant epithelial cells, glandular mucus cells, and endothelial cells. This single-cell level expression pattern provides a biological explanation for the weak correlation between MET and immune cell infiltration observed in large sample transcriptome data, further supporting the conclusion that MET mainly exerts carcinogenic effects through the intrinsic program of tumor cells rather than directly regulating the immune microenvironment.

**Fig. 8.**
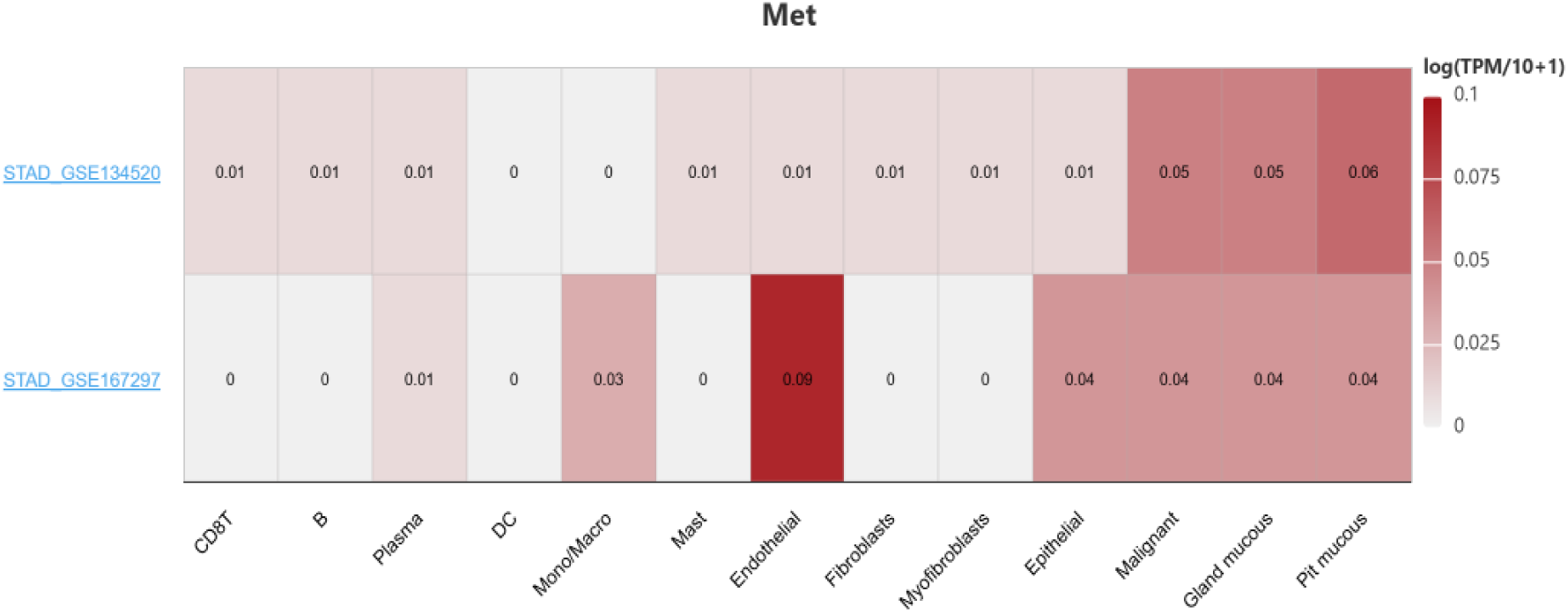
Cell-type-specific expression landscape of MET in two independent gastric cancer single-cell RNA-seq cohorts (GSE134520 and GSE167297). The color gradient corresponds to the normalized expression value log(TPM/10+1) of MET, with deeper red indicating higher expression.

### 2.9 Correlation analysis between MET and core immune checkpoint molecules

To further improve immune related analysis, we utilized the GEPIA2 database to investigate the correlation between MET expression and key immune checkpoint molecules (Figure 9A-F). The results showed that MET expression was not significantly correlated with PD-1 (r=-0.05, *P*= .31, Figure 9A), PD-L1 (r=-0.0036, *P*= .94, Figure 9B), CTLA4 (r=-0.026, *P*= .60, Figure 9C), TIGIT (r=-0.027, *P*= .59, Figure 9D), LAG3 (r=-0.019, *P*= .71, Figure 9E), and HAVCR2 (r=0.067, *P*= .15, Figure 9F). The absolute correlation coefficients of all paired combinations were all below 0.1, with no statistical significance, indicating that MET expression levels do not directly regulate the transcription levels of immune checkpoint molecules such as PD-1/PD-L1, CTLA4, TIGIT, LAG3 and HAVCR2.

**Figure 9.**
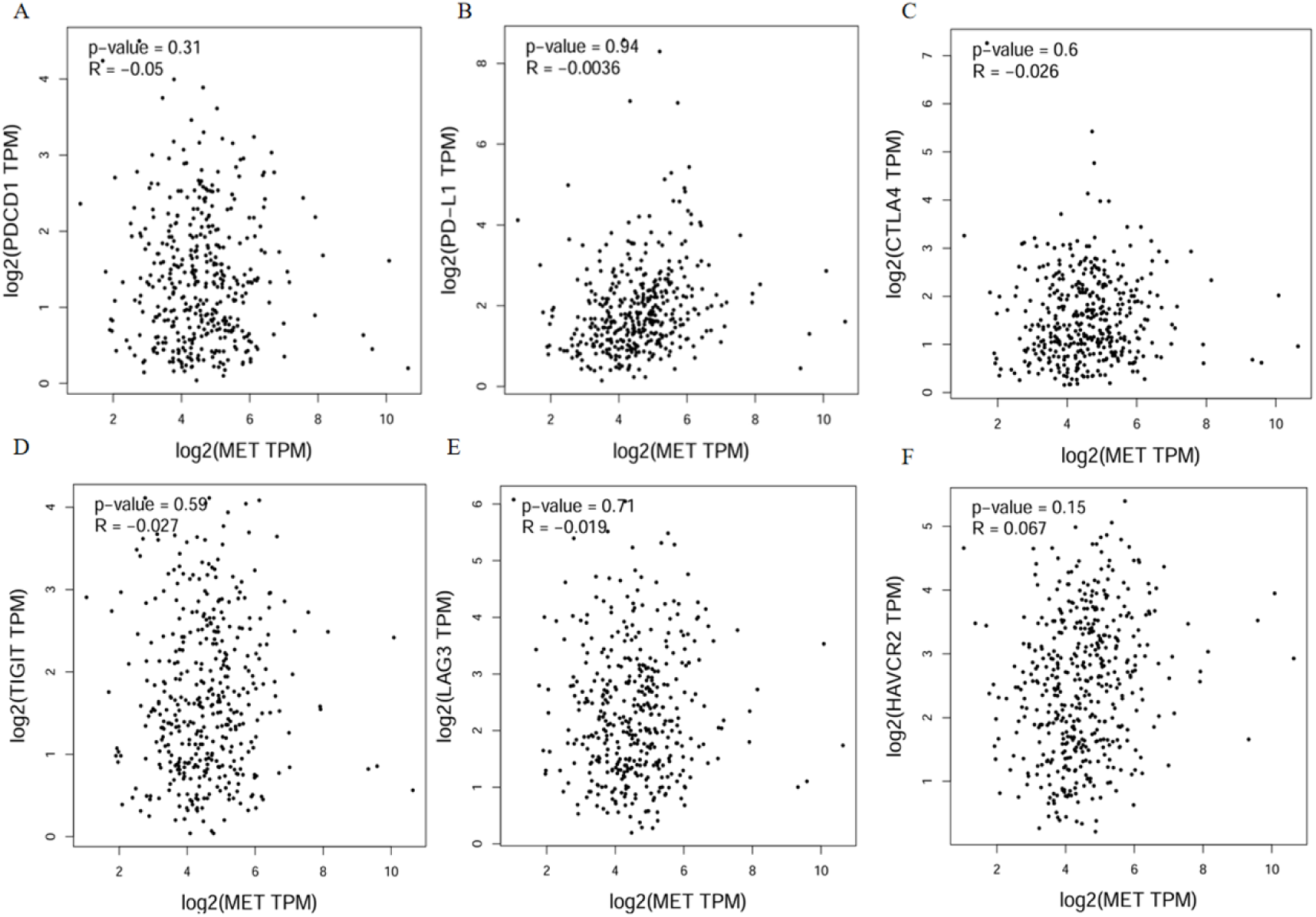
Correlation scatter plots between MET and six core immune checkpoint genes in TCGA-STAD cohort generated by GEPIA2. X-axis represents log₂-transformed TPM expression of MET, Y-axis represents log₂-transformed TPM expression of immune checkpoint genes. A: PDCD1 (PD-1); B: PD-L1; C: CTLA4; D: TIGIT; E: LAG3; F: HAVCR2.

## 3 Discussions

MET is a proto-oncogene encoding a tyrosine kinase receptor located on chromosome 7 (7q21-q31), with hepatocyte growth factor (HGF) as its ligand[13, 14]. Previous studies have demonstrated that MET is closely associated with various malignancies, including lung cancer[15], liver cancer[16] and gastric cancer[17], where it promotes tumor proliferation, invasion, metastasis, and angiogenesis[18, 19]. Currently, MET tyrosine kinase inhibitors (TKIs) such as savolitinib and capmatinib have been widely used in the treatment of advanced non-small cell lung cancer[20, 21]. However, the clinical significance of MET targeting in gastric cancer remains controversial. Therefore, elucidating the specific role of MET in gastric cancer development is of great importance for guiding targeted therapy and prognosis evaluation.

This study systematically depicted the expression characteristics, prognostic value, molecular regulatory network, and immune infiltration pattern of MET in gastric cancer through bioinformatics analysis of multiple data integrations. The study found that the expression level of MET in gastric cancer tissue was significantly higher than that in normal gastric mucosal tissue, and its high expression was associated with tumor differentiation and TNM staging, which is consistent with the previous research results of Metzger et al.[22]. Further analysis shows that there is no clear correlation between MET expression levels and clinical pathological characteristics such as gender and age in gastric cancer patients, suggesting that MET overexpression may be an early or universal event in the development of gastric cancer, rather than a late stage tumor specific change. The survival analysis results showed that the overall survival and progression free survival of patients with high MET expression were significantly lower than those with low MET expression, which is consistent with the meta-analysis results of Peng et al.[23], which showed that the mortality rate of gastric cancer patients with high MET expression and MET amplification was significantly increased. We also found that MET is also highly expressed in cervical squamous cell carcinoma, colon adenocarcinoma and other cancers, while its expression is reduced in breast cancer, acute myeloid leukemia and low-grade glioma, suggesting that the expression pattern of MET is cancer specific.

Protein interaction analysis showed that MET has direct or indirect interactions with multiple HER family receptor tyrosine kinases such as EGFR and ERBB2. This finding provides a potential explanation for the poor efficacy of anti-HER-2 therapy in clinical practice: when ERBB2 is inhibited, MET signaling may be activated as an "escape pathway". Meanwhile, tumor cells may also activate other pathways such as RTK to bypass MET inhibition and develop resistance to MET inhibitors[24]. Therefore, for gastric cancer patients with high MET expression, multi-target combined inhibition may be a more effective treatment strategy than monotherapy.

Functional enrichment analysis showed that at the level of GO biological processes, differentially expressed genes were significantly enriched in the ERBB signaling pathway, integrin mediated cell adhesion, and other processes, suggesting that MET may affect the malignant phenotype of gastric cancer cells by regulating the receptor tyrosine kinase signaling network and cell adhesion ability. At the hierarchical level of GO cells, entries such as DNA repair complexes and chromatin related structures were significantly enriched, providing subcellular localization evidence for MET’s involvement in genome stability regulation. At the functional level of GO molecules, the binding function of calcium binding proteins is significantly enriched, while the functions related to oxidoreductases only show an enrichment trend and have not reached statistical significance.

More importantly, KEGG pathway enrichment analysis showed that MET related genes were significantly enriched in DNA damage repair pathways such as homologous recombination repair and mismatch repair. Homologous recombination repair is the core mechanism for cell repair of DNA double strand breaks. Tumor cells with normal homologous recombination repair function can efficiently repair platinum induced DNA damage, leading to chemotherapy resistance[25]. Based on the extensive interactions observed between MET and RTK family members such as EGFR and ERBB2 in the PPI network, we infer that gastric cancer patients with high MET expression may face a dual dilemma of "chemotherapy resistance caused by homologous recombination repair activation" and "targeted monotherapy resistance caused by RTK cross activation". It suggests that MET is not only a prognostic marker for gastric cancer, but also a potential indicator for predicting chemotherapy sensitivity. For gastric cancer patients with high MET expression, the efficacy of conventional platinum based chemotherapy may be limited, and it is necessary to consider adjusting the chemotherapy regimen or combining targeted therapy to overcome drug resistance.

In addition, KEGG analysis also found a significant enrichment of bacterial invasion pathways into epithelial cells. However, considering the weak direct correlation between this pathway and MET’s pro cancer mechanism, this study did not focus on it as a key analysis direction. In addition to the core pathways mentioned above, the enrichment of nuclear cytoplasmic transport, oxidative phosphorylation and other pathways also suggests that the carcinogenic effect of MET involves multiple levels such as transcriptional regulation and energy metabolism reprogramming, which together form a complex and collaborative pro cancer network.

In terms of tumor immune microenvironment, this study systematically evaluated the relationship between MET and immune cell infiltration through cross validation using four independent algorithms: TIMER, CIBERSORT, EPIC, and MCPcounter. The results showed a robust negative correlation between MET and B cells infiltration (consistent across all four algorithms), and algorithmic heterogeneity in its association with neutrophils, CD8 ⁺T cells, CD4 ⁺T cells, and macrophages. To further validate this result, we supplemented the correlation analysis between MET and core immune checkpoint molecules and found no significant association between MET and PD-1, PD-L1, CTLA4, TIGIT, LAG3, HAVCR2. In addition, two independent single-cell external queue data also confirmed that MET is almost not expressed in various immune cells such as CD8 ⁺ T cells, B cells, dendritic cells, macrophages, etc. Its transcripts are mainly enriched in malignant epithelial cells and endothelial cells. These findings provide preliminary clues, suggesting that MET does not directly regulate anti-tumor immunity in gastric cancer, and its oncogenic effect mainly depends on the inherent signaling pathways of tumor cells. The finding that MET is not significantly correlated with immune checkpoint molecules suggests that the combined benefits of MET targeted drugs and immunotherapy may not be achieved through direct inhibition of immune checkpoint pathways. Of course, in the absence of functional validation evidence, this result is an exploratory discovery rather than a definitive conclusion.

Although multiple layers of evidence confirm that MET has limited direct regulatory effects on the immune microenvironment of gastric cancer, the negative correlation between MET and B cell infiltration observed across four algorithms in this study still has certain reference value. In vitro and animal experiments have shown that the c-MET signaling pathway can inhibit the body’s anti-tumor immune respons[26]. However, based on the single-cell sequencing results of this study, MET is almost not expressed in immune cells such as B cells, indicating that the negative correlation between the two is only an indirect effect of tumor microenvironment remodeling, rather than the inhibitory effect of MET directly acting on B cells. Overall, the weak association between MET and tumor immune cell infiltration is only an exploratory clue, and the exact molecular mechanism of their interaction still needs further verification through functional experiments.

This study further supplemented the correlation analysis between MET and core immune checkpoint molecules, and the results showed that MET was not significantly associated with PD-1, PD-L1, CTLA4, TIGIT, LAG3, and HAVCR2. The above multiple layers of evidence collectively confirm that MET does not have the function of directly regulating anti-tumor immunity in gastric cancer, and its pro cancer effect mainly depends on the inherent signaling pathways of tumor cells (RTK cross activation, homologous recombination repair abnormalities, etc.). The finding that MET is not significantly correlated with immune checkpoint molecules has clinical reference value. The results of this study do not support the direct upregulation of immune suppressive molecules such as PD-L1 by MET, indicating that the combined benefits of MET targeted drugs and immunotherapy cannot be achieved through direct inhibition of immune checkpoint pathways. The combination strategy of the two requires clinical exploration based on other indirect mechanisms.

This study also has certain limitations. Firstly, all analyses are based on batch transcriptome sequencing data provided by public databases, which may be influenced by factors such as sequencing platform differences, tumor heterogeneity, and sample purity[27]; As a highly heterogeneous tumor, gastric cancer may have mixed transcriptional information of cancer cells, stromal cells, and immune cells in the sample, which may dilute or distort the expression signal of MET. Therefore, the observed differences and correlations in gene expression may not fully reflect the true molecular state of tumor cells. Secondly, this study is a bioinformatics correlation analysis, and the hypotheses proposed for RTK cross activation, homologous recombination repair pathway abnormalities, and immune microenvironment regulation are all computational predictions. The exact causal relationship and molecular mechanism still need further verification through in vitro and in vivo functional experiments. In addition, third, this study mainly explored the expression and function of MET at the transcriptome level, without including cross validation at multiple omics levels such as proteomics, metabolomics, imaging, and spatial transcriptomics. Finally, this study lacks independent prospective clinical cohort validation, and the predictive value of MET for chemotherapy and targeted therapy efficacy still needs further evaluation in clinical practice.

Future research can integrate PET/CT imaging[28], single-cell sequencing, spatial transcriptomics, and multi omics data, rely on biosensor platforms to achieve real-time dynamic monitoring of MET related biomarkers[29], and combine potential biomarkers such as MET with personalized vaccine development strategies[30]. Through cell and animal experiments, the regulatory effects of MET on RTK signal crosstalk, homologous recombination repair pathways, immune microenvironment, and immune checkpoint molecules were systematically validated. By collecting clinical gastric cancer cohorts, we aim to verify the relationship between MET expression and platinum based chemotherapy, as well as the correlation with targeted drug efficacy. We aim to explore the anti-tumor effects of MET inhibitors in combination with EGFR/ERBB2 inhibitors, PARP inhibitors, and other drugs, providing experimental evidence for evaluating their value as efficacy prediction markers and related therapeutic drugs. Through the above multi-level research, it is expected to more comprehensively reveal the mechanism of action of MET in gastric cancer, providing new targets and strategies for precise treatment of gastric cancer.

## 4 Conclusion

In summary, this bioinformatic study systematically characterized MET expression, prognostic value and molecular regulatory network in gastric cancer across multiple databases. Our analyses show that MET is markedly upregulated in gastric cancer and correlates with advanced tumor grade, late TNM stage and poor survival, supporting its potential as a prognostic biomarker. MET interacts with 20 key signaling molecules including EGFR, ERBB2, HGF, STAT3 and GRB2. Functional enrichment reveals enrichment in ERBB signaling, cadherin binding and DNA repair complexes, especially homologous recombination and mismatch repair pathways. These findings suggest that MET overexpression may trigger RTK compensatory activation and enhanced DNA repair capacity, which could lead to dual resistance to targeted monotherapy and chemotherapy, providing a rationale for combination treatment strategies. Multi-algorithm immune infiltration analyses combined with single-cell data demonstrate only weak, algorithm-dependent associations between MET and immune cell infiltration. Together with the observation of negligible MET expression in immune cells, these results indicate MET exerts limited direct regulation on the gastric cancer immune microenvironment, with its oncogenic effects mainly mediated via intrinsic tumor-cell signaling. Collectively, this work provides multidimensional bioinformatic evidence for MET as a candidate prognostic biomarker and therapeutic target in gastric cancer. Further functional experiments and prospective cohorts are required to validate this mechanistic hypothesis.

## Acknowledgments

Not applicable

## Funding

Wuwei Science and Technology Bureau (project number WW24B01SF087).

2026 Digestive Medicine Gansu Province Clinical Key Specialty Capability Enhancement Project.

## Author contributions

Zhengjie Zhang and Xiuzhen Wei designed and drafted manuscript.

## Institutionan Review Board Statement

Not applicable

## Informed Consent Statement

Not applicable

## Consent to publication

All authors rare approved the final manuscript and the submission to this journal.

## References

[1]. Siegel, R.L., et al., Cancer statistics, 2026. CA Cancer J Clin, 2026. 76(1): p. e70043.

[2]. López, M.J., et al., Characteristics of gastric cancer around the world. Crit Rev Oncol Hematol, 2023. 181: p. 103841.

[3]. Ajani, J.A., et al., Gastric Cancer, Version 2.2025, NCCN Clinical Practice Guidelines In Oncology. J Natl Compr Canc Netw, 2025. 23(5): p. 169–191.

[4]. Sundar, R., et al., Gastric cancer. Lancet, 2025. 405(10494): p. 2087–2102.

[5]. Patel, A.K., N.S. Sethi and H. Park, Gastric Cancer: A Review. JAMA, 2026. 335(5): p. 439–450.

[6]. Bang, Y., et al., Trastuzumab in combination with chemotherapy versus chemotherapy alone for treatment of HER2-positive advanced gastric or gastro-oesophageal junction cancer (ToGA): a phase 3, open-label, randomised controlled trial. Lancet, 2010. 376(9742): p. 687-97.

[7]. Shah, M.A., et al., Zolbetuximab plus CAPOX in CLDN18.2-positive gastric or gastroesophageal junction adenocarcinoma: the randomized, phase 3 GLOW trial. Nat Med, 2023. 29(8): p. 2133–2141.

[8]. Chen, W., et al., Trastuzumab Deruxtecan Resistance via Loss of HER2 Expression and Binding. Cancer Discov, 2026. 16(2): p. 235-249.

[9]. Guo, R., et al., MET-dependent solid tumours - molecular diagnosis and targeted therapy. Nat Rev Clin Oncol, 2020. 17(9): p. 569–587.

[10]. Röcken, C., Predictive biomarkers in gastric cancer. J Cancer Res Clin Oncol, 2023. 149(1): p. 467–481.

[11]. Kuniyasu, H., et al., Frequent amplification of the c-met gene in scirrhous type stomach cancer. Biochem Biophys Res Commun, 1992. 189(1): p. 227–32.

[12]. Ahmad Alizadeh, E., et al., Elucidating the Inhibitory Potential of Statins Against Oncogenic c-Met Tyrosine Kinase Through Computational and Cell-based Studies. Iran J Pharm Res, 2025. 24(1): p. e158845.

[13]. Duh, F.M., et al., Gene structure of the human MET proto-oncogene. Oncogene, 1997. 15(13): p. 1583–6.

[14]. Dean, M., et al., The human met oncogene is related to the tyrosine kinase oncogenes. Nature, 1985. 318(6044): p. 385-8.

[15]. Remon, J., et al., MET alterations in NSCLC-Current Perspectives and Future Challenges. J Thorac Oncol, 2023. 18(4): p. 419–435.

[16]. Wang, T., et al., MET promotes hepatocellular carcinoma development through the promotion of TRIB3-mediated FOXO1 degradation. Clin Mol Hepatol, 2025. 31(3): p. 1032–1057.

[17]. Zhang, Y., L. Shen and Z. Peng, Advances in MET tyrosine kinase inhibitors in gastric cancer. Cancer Biol Med, 2024. 21(6): p. 484–98.

[18]. Gallo, S., C.B. Folco and T. Crepaldi, The MET Oncogene Network of Interacting Cell Surface Proteins. Int J Mol Sci, 2024. 25(24).

[19]. Hyung, S., et al., Patient-derived exosomes facilitate therapeutic targeting of oncogenic MET in advanced gastric cancer. Sci Adv, 2023. 9(47): p. eadk1098.

[20]. Yu, Y., et al., Savolitinib in patients in China with locally advanced or metastatic treatment-naive non-small-cell lung cancer harbouring MET exon 14 skipping mutations: results from a single-arm, multicohort, multicentre, open-label, phase 3b confirmatory study. Lancet Respir Med, 2024. 12(12): p. 958–966.

[21]. Wolf, J., et al., Capmatinib in MET Exon 14-Mutated or MET-Amplified Non-Small-Cell Lung Cancer. N Engl J Med, 2020. 383(10): p. 944–957.

[22]. Metzger, M., et al., MET in gastric cancer--discarding a 10% cutoff rule. Histopathology, 2016. 68(2): p. 241–53.

[23]. Peng, Z., et al., Prognostic significance of MET amplification and expression in gastric cancer: a systematic review with meta-analysis. PLoS One, 2014. 9(1): p. e84502.

[24]. Yao, S., et al., Unveiling the Role of HGF/c-Met Signaling in Non-Small Cell Lung Cancer Tumor Microenvironment. Int J Mol Sci, 2024. 25(16).

[25]. Shah, B., M. Hussain and A. Seth, Homologous Recombination Deficiency in Ovarian and Breast Cancers: Biomarkers, Diagnosis, and Treatment. Curr Issues Mol Biol, 2025. 47(8).

[26]. Koh, S.A. and K.H. Lee, Function of hepatocyte growth factor in gastric cancer proliferation and invasion. Yeungnam Univ J Med, 2020. 37(2): p. 73–78.

[27]. Liu, H., et al., Technical and Biological Biases in Bulk Transcriptomic Data Mining for Cancer Research. J Cancer, 2025. 16(1): p. 34–43.

[28]. Singh, S.B., et al., PET/CT in the Evaluation of CAR-T Cell Immunotherapy in Hematological Malignancies. Mol Imaging, 2024. 23: p. 15353508241257924.

[29]. Nimhan, G., M. Narwade and K. Gajbhiye, Biosensor driven biomarker analysis: pioneering advancements in cancer diagnosis and therapeutic strategies. Biomarkers, 2025. 30(4): p. 332–351.

[30]. Iamukova, L. and E. Alferova, Personalized Cancer Vaccines in the Clinical Trial Pipeline. Asia Pac J Clin Oncol, 2026. 22(3): p. 362-368.

